# Role of aridity in shaping adaptive genomic divergence and population connectivity in a Southern African rodent

**DOI:** 10.1101/2025.07.29.667317

**Authors:** Hamilcar Socrates Keilani, Anna-Sophie Fiston-Lavier, Pierre Caminade, Anne Loiseau, Maxime Galan, Hugues Parrinello, Carine Brouat, Raphael Leblois, Nico Avenant, Neville Pillay, Guila Ganem, Carole M. Smadja

## Abstract

Elucidating the drivers of evolution in dry environments is central to understanding how organisms respond to climate change. While research on the genomics of adaptation is growing, aridity-driven intraspecific divergence remains poorly quantified. Here, we address this gap by using genomic data from 230 individuals of the arid-adapted four-striped mouse *Rhabdomys bechuanae*, sampled across an aridity gradient in southern Africa, a region facing increasing aridification. Combining these data with palaeoclimatic reconstructions and present-day aridity indices, we investigate, from a spatio-temporal perspective, how intraspecific genetic variation relates to aridity. Inference of past effective population size revealed a sharp decline in the late Pleistocene, coinciding with regional aridification and potentially reflecting changes in connectivity during dry periods. Current population structure followed a pattern of isolation by distance and mirrored the aridity gradient. Genotype-Environment Association analyses identified SNPs and genes significantly associated with aridity and genetically differentiated among populations, with functions related to water and energy conservation - as expected under arid conditions - as well as neurotransmission. These findings highlight the underappreciated role for neurological processes in coping with water and resource scarcity. More broadly, our integrative genomics approach suggests that aridity shapes population connectivity and adaptation, with implications for climate resilience.

## Introduction

Climate change has increasingly been associated with local extinctions [1], shifts in phenology [2] and alteration of species’ geographic distributions [3]. These responses tend to be particularly pronounced in harsh environments, such as arid regions, where species already face several ecological and physiological constraints [4]. With projections indicating a rise in the intensity and frequency of drought events in the coming decades [5,6], it is essential to better understand how species respond to both temporal and spatial variation in aridity. By combining newly generated genomic data with available palaeoclimatic reconstructions and contemporary measures of aridity, this study aims to shed light on the role of aridification on the demographic history, spatial population structure and adaptive responses of a non-model rodent species. These insights are key for predicting resilience of natural populations under future climate scenarios and for guiding conservation strategies [7].

At the individual level, the primary challenges of life in arid environments include maintaining body temperature and conserving water [8]. These challenges are especially evident for species that rely on evaporative water loss mechanisms, such as sweating, to dissipate heat, particularly in mammals [9]. The physiological demands imposed by such environmental conditions, particularly when experienced over multiple generations, often exceed the limits of phenotypic plasticity in non-desert-adapted species. Consequently, it is assumed that strong selection has acted on complex physiological traits related to energy metabolism, water retention and thermoregulation in species inhabiting arid environments [8–10].

Genomic studies have identified recurrent genetic patterns among mammals living in deserts [10]. Evidence of adaptation to arid environments has included elevated allele frequency differentiation [11,12], reduced genetic diversity [11,13,14], increased non-synonymous-to-synonymous substitution rate ratios [15] and divergent gene expression patterns [16] in genomic regions involved in key functions for desert adaptation. These genomic regions often relate to key physiological traits relevant to the survival of both mammalian specialist and non-specialist species in deserts, including DNA repair, protein synthesis and degradation [14,15], thyroid hormone levels [11], water retention [17], lipid [15] and carbohydrate metabolism [11,13,18,19]. However, most of these studies have relied on qualitative comparisons between desert and non-desert species or populations, and relatively few have explicitly explored quantitative variation along continuous environmental gradients. Several other studies on the genomics of local adaptation have demonstrated the power of Genotype-Environment Association (GEA) analyses to identify polymorphisms associated with continuous environmental variation [20,21]. In particular, Rocha et al. [10] highlighted the use of the Aridity Index (AI), known to be a good predictor of certain aridity-adapted phenotypes, as a powerful, composite and quantitative environmental variable to inform genomic studies of adaptation to arid conditions. Aridity Index has been shown to be a major driver of ecosystem structure and functioning in drylands, and exhibits important ecological shifts along gradients of aridity [22].

*R. bechuanae*, a four-striped mouse species from Southern Africa, is a good model species to study quantitative variation across gradients and GEAs, as its range overlaps with hot semi-arid to hyper-arid summer rainfall regions of Southern Africa [23]. Divergence from its closest relatives within the *Rhabdomys* species complex occurred approximately 3.1-4.3 million years ago [24], and this diversification has been associated with environmental niche differentiation [25]. It likely reflects a long-term trend towards increasing aridity and seasonality in Southern Africa since the late Miocene, coupled with a shift from closed subtropical woodland to sparse, shrubby vegetation [26]. Phenotypic traits suggest adaptation to aridity; these traits include morphological (longer tail length related to potential differences in locomotion and/or thermoregulation) and behavioural divergence (selection of different habitats) from more mesic congeners [21,27], as well as enhanced osmoregulatory ability, as suggested by differences in blood metabolite concentrations between populations of *R. bechuanae* and of its sister species *R. dilectus,* occurring syntopically in semi-arid environments [28]. In this study, we hypothesised that this aridity gradient would impose divergent selective pressures on key physiological traits related to adaptation to aridity, leading to detectable intraspecific population differentiation and genetic variation at loci associated with aridity-related functions.

To investigate genomic signatures of evolution under arid conditions across an extensive sample of individuals (n=230) from multiple localities spanning the distribution of *R. bechuanae* along an aridity gradient, we generated a genome-wide SNP dataset using restriction-site associated DNA sequencing (RAD-seq) [29], a reduced-representation method adapted to generating population-level genomic data in non-model species with large genome sizes [30]. Indeed, our overarching aim was to evaluate how the temporal and spatial variation in aridity has shaped the population history and structure of *R. bechuanae*, by integrating inferences of historical changes in effective population size with GEA analyses.

Our first objective was to reconstruct past changes in effective population size in relation to palaeoclimatically inferred shifts in aridity. We expected that historical episodes of increased aridity would be associated with population declines, either due to demographic bottlenecks [31,32], or via reductions in genetic diversity caused by habitat fragmentation and decreased population connectivity. As spatial variation in aridity may also influence current population structure and genetic diversity [12,33], our second objective was to characterise the current population structure within *R. bechuanae* and assess its relationship with spatial variation in aridity across the species’ range. We predicted that population structure would correlate with the degree of aridity, reflecting its impact on gene flow and differentiation. Our third objective was to assess the genomic basis of adaptation to aridity in *R. bechuanae* using a GEA approach with the Aridity Index as a quantitative predictor. This approach enabled us to identify loci associated with spatial variation in arid conditions across the species’ range and to identify potential biological functions that may underpin survival in arid environments. We hypothesised that site frequency variation would concentrate in genomic regions linked to key functions for survival in deserts such as DNA repair, protein modification, osmoregulation, and carbohydrate/fat metabolism [10,15].

## Material & Methods

### Samples, aridity index and RAD-seq data production

Individual samples of four-striped mice of the genus *Rhabdomys* were previously collected in their natural environment between 2007 and 2019 by us or collaborators (details for trapping procedures in [23]). Based on data obtained for *R. pumilio* indicating minimal relatedness at this distance [34], we aimed to avoid related individuals as much as possible, by selecting mice trapped at least 100 m apart, except for breeding pairs (a male and a female).

To study fine-scale intraspecific structure, as well as perform GEA and historical effective population size inferences in *R. bechuanae*, we selected a large number of individuals from this species, sampled in 11 different localities at least 20 km distant from each other and along the aridity gradient (*N*=230 individual samples in total, 19.82 +/- 4.72 individuals per locality, **Table S1**). We estimated the aridity level of each *R. bechuanae* sampling locality by computing the Aridity Index (AI) at each location, following [35]] (**Supplementary Material** for details). Species identification for all sampled individuals in the *Rhabdomys* species complex composed of cryptic species had previously been obtained via COI sequencing or RFLP genotyping ([23] and procedure therein; **Table S1**). DNA extraction, RAD-sequencing library preparation and Ilumina paired-end (2x150bp) sequencing procedures are described in **Supplementary Material.**

### RAD-seq raw data processing and filtering

Duplicate reads resulting from PCR amplification were discarded using the program *clone_filter* implemented in the *Stacks* v. 2.66 software [36]. The resulting paired-end reads were demultiplexed and quality filtered using the *process_radtags* pipeline, also implemented within *Stacks*. Sequence reads were aligned to the *R. pumilio* reference genome (2.3 Gb in length; GCA_030674055.1, [37]), using *BOWTIE* (v.2.4.5) [38]. Before mapping, putative repeated elements were hard-masked (i.e., repeated sequences were replaced by stretch of “N”) from the reference genome using the mouse repeated element Repbase Update (v.27.06) database using *Repeatmasker* v.4.1.3. [39]. Reads mapping to more than one reference sequence were discarded, and the maximum number of mismatches allowed was three. The resulting files were converted to .bam files and sorted using *SAMtools* v.1.18 [40].

The reference-based *Gstacks* pipeline was applied to cluster reads into RAD loci using the Marukilow model, minimum mapping quality of 40, and alpha thresholds (for mean and variance) of 0.05 for discovering single nucleotide polymorphisms (SNPs). Using the catalogue of loci generated with *Gstacks*, the *populations* program was then used to proceed to SNP and sample filtering steps and generate SNP datasets adapted to each type of downstream analyses (**Supplementary Material** and **Table S2** for details).

### Inferences of historical changes in effective population size

To evaluate historical events that could have significantly impacted current population genetic diversity and structure, and test whether historical variations in effective population size (*N_e_*) coincide with known past variations in aridity, we estimated historical changes in *N_e_* using *Stairway Plot 2* v.2.1.2, which conducts multi-epoch coalescent inference to infer changes in population size through time [41,42]. We performed this analysis at two different sampling scales: 1) at the *R. bechuanae* species level, pooling individual samples from all sampled localities, and 2) at the locality level, analysing two localities representing the two extremes of the Aridity Index across the sampled distribution.

Because site frequency spectrum (SFS)-based demographic inference is particularly sensitive to rare alleles, especially singletons and doubletons [43], we used the SNP dataset obtained without applying a minor allele count (-mac) filter. To minimise the effects of missing data while retaining a sufficient number of SNPs for historical effective population size inference, we implemented two complementary filtering steps. First, to homogenise sampling effort and avoid biases due to unequal sample sizes, we applied a downsampling procedure with a maximum of 19 individuals per locality. During this step, individuals with the highest proportions of missing genotypes were preferentially excluded, thereby reducing overall missing data in the final datasets. Second, folded SFS were generated using *easySFS* [44], which implements a projection approach that retains loci only when sufficient genotype information is available, maximising SNP inclusion while limiting biases from uneven missingness among samples. Filtering outcomes - retained number of individuals and SNPs, mean depth of coverage per site, and mean missing data per site for each final projected dataset - are provided in Table **S3**.

For all *Stairway Plot 2* analyses (see **Supplementary Material** for details), the mutation rate was set to 5.7 × 10^−9^ per site per generation (based on the germline mutation rate estimated in *Mus musculus* in [45]), and the generation time was assumed to be one year. We considered the entire allele frequency spectrum. However, we performed a sensitivity analysis after excluding singleton variants to test their effect on *Stairway Plot 2* inference.

### Population structure analyses

For these analyses, we also applied a downsampling procedure to the filtered SNP dataset with a maximum of 19 individuals per locality, selecting individuals with the least missing data. To describe the structure of genetic diversity within *R. bechuanae*, we first computed differentiation statistics (pairwise *F_ST_*) [46] for all sampled localities using the *vcftools* program v.0.1.16 [47]. To test isolation-by-distance and isolation-by- environment within *R. bechuanae*, simple and partial Mantel tests (R *ncf* package v.1.3-2) [48] were conducted to test for correlations between genetic differentiation among all *R. bechuanae* localities (using *F_ST_*/(1- *F_ST_)* following [49]) and log-transformed geographic distance, as well as log-transformed environmental distance (using the Aridity Index). To characterise population structure within *R. bechuanae*, we performed a principal component analysis (PCA) and a sparse non-negative matrix factorisation (*sNMF)* analysis using the R package *LEA* v.3.14 [50]. For *sNMF* analyses, we ran 1,500 repetitions for each value of *K* ranging from 1 to 8 and chose the best *K* by evaluating cross-entropy values. Additionally, a neighbour-joining tree was generated using SNP data of these same samples with the R package *ape* (v.5.7-1) [51].

### Genotype-Environment Association analyses

To explore signals of genetic adaptation to aridity, we conducted Genotype–Environment Association (GEA) analyses. To test for an association between allele frequency differences between all *R. bechuanae* localities and the environmental variable of interest, the Aridity Index, which was computed for all localities using [35], we used the program *BayPass* v.2.4 [22]. We first pooled individual genotype information from each of the 11 *R. bechuanae* localities, before running *BayPass* under the standard covariate model, which uses an Importance Sampling (IS) approximation to identify SNPs associated with the Aridity Index, and corrects for population structure by using the scaled covariance matrix Ω of population allele frequencies. Three runs (with different starting seeds) were performed to test for convergence. To determine the level of association of each SNP with the Aridity Index and identify SNPs significantly associated, we computed the empirical Bayesian p-value (eBPis) and Bayes factors (*BF*) expressed in deciban units (dB), with 10 dB being a threshold of strong association according to Jeffreys’ rule [52]. Additionally, *XtX* statistics, which explicitly account for population structure through Ω, were outputted by the *BayPass* core model to identify SNPs with significant genetic differentiation among *R. bechuanae* populations (**Supplementary Material** and **Figure S1** for details).

To construct an accurate gene annotation file, we mapped the *Mus musculus* GRCm39 assembly annotation [53] onto the *R. pumilio* genome using the default settings of *liftoff* software version 1.6.3 [54]. All SNPs were mapped onto the annotation, and we identified all annotated protein-coding genes containing genotyped SNPs for functional analyses. Among those, we defined genes containing SNPs that were found to be significantly associated with the Aridity Index (*BF* greater than 10 dB) were defined as outlier ‘association’ genes, and among those, genes containing SNPs both significantly associated and differentiated among populations (both with a *BF* greater than 10 dB and a significant *XtX*) as outlier ‘association/differentiation’ genes. For each outlier gene, we first retrieved a full description from the Alliance of Genome Resources [55], which was then summarised to obtain a meaningful set of functional information. We performed functional enrichment analyses on these two lists of outlier genes using *Metascape*, an online tool for gene function annotation analysis. Gene Set Enrichment Analysis (GSEA) is a computational method that determines whether an *a priori* defined set of genes contains a statistically significant overrepresentation of biological functions or states. Here, Gene Ontology (GO) terms, a standardised framework to describe the roles of genes and their products, were considered. The enrichment was initially considered statistically significant at *p* < 0.05. P-values were then corrected for false discovery rates according to the Benjamini-Hochberg method. Only enrichments exhibiting a corrected *p_ad_*_j_ < 0.05 were considered significant.

## Results

### RAD-seq and genotype datasets, Aridity Index estimates

Across all 230 *R. bechuanae* samples analysed, an average of 2,316,057 sequence reads per sample were mapped against the *R. pumilio* reference genome, after using *process_radtags* to filter for quality and *clone-filter* to mask potential PCR duplicates (see **Table S4** for details). The alignment rate to the repeat-masked *R. pumilio* genome averaged 55.96 % across *R. bechuanae* samples. The *Gstacks* analysis resulted in an average of 151,936 ± 1,779 loci per sample. RAD loci were genotyped with a mean per-sample coverage of 10.94 ± 0.21 and a mean of 227.88 ± 0.09 base pairs per locus.

The SNP and sample filtering procedures used to generate the three different *R. bechuanae* datasets for population structure analyses, past effective population size inference and GEA analyses resulted in final datasets comprising 44,900 SNPs for the population structure and GEA analyses, and more than 145,000 SNPs for past effective population size at both the species and locality levels (see **Tables S2** and **S3** for details). A total of 152 samples were retained for population structure and historical population effective size analyses, and 218 samples for the GEA analyses (**Tables S2, S3, S4**). After all filtering steps, the three final datasets exhibited mean per-sample coverages of 18.39 ± 0.43, 18.06 ± 0.43, and 18.02 ± 0.38, respectively, and mean proportions of missing data per sample of 10.65 ± 0.79, 7.09 ± 0.59, and 11.99 ± 0.72 % (**Table S4**). The additional filtering steps for SFS-based *Stairway Plot 2* analyses retained a large number of SNPs while maintaining very low levels of mean missing data per site. Mean data missingness ranged from 3% to 8% across the three datasets analysed (Klein Pella, Sandveld and species-level datasets; **Table S3**). After *easySFS* projection, 258 projected chromosomes (N=129 samples) were retained for species-level *Stairway Plot 2* analyses; 28 and 32 projected chromosomes were retained for Klein Pella and Sandveld analyses, respectively.

Aridity Index (AI) estimations validated our sampling of *R. bechuanae* along an aridity gradient in South Africa: AI of sampled localities ranged from 0.0322 (Klein Pella, classified as hyper-arid) to 0.2457 (Sandveld, classified as semi-arid) (**Table S5**).

### Genetic diversity and population structure in time and space

Historical changes in effective population size in *R. bechuanae*

Folded site frequency spectra (SFS) obtained after all filtering steps and *easySFS* projection are shown in **Figure S2**. For the species-level dataset, the frequency of singleton variants was lower than that of doubletons, indicating a deficit in singleton alleles relative to expectations under standard neutral models. In contrast, the two locality-level datasets showed SFS distributions more consistent with the expected decline in allele frequencies across allelic classes. To evaluate whether this deficit in singleton variants influenced *Stairway Plot 2* inference for the species-level dataset, we repeated this analysis after excluding singletons from the SFS. The inferred trajectory of historical change in *N_e_* at the species-level was qualitatively very similar to those obtained using the complete SFS (**Figure S3**), indicating that the demographic signal was primarily carried by the remaining allele frequency classes rather than by singleton variants. This pattern supports the robustness of our inference despite the apparent depletion of singletons. We detected no major differences in the temporal patterns of *N_e_* between the two populations located at the semi-arid (Sandveld) or hyper-arid (Klein Pella) extremes of the aridity gradient, nor any evidence of recent population decline (**Figure 1**, green and red thin lines). Both populations showed relatively stable and similar *N_e_* trajectories at least from the Last Glacial Maximum (LGM) to the present. At the species level, pooling individuals from all sampled localities resulted in a markedly different demographic trajectory (**Figure 1**, thick red line). Similar to population-specific analyses, *Stairway Plot 2* inferred a strong increase in *N_e_* beginning approximately 140 thousand years ago (kya), followed by stabilisation around 100 kya, with a median *N_e_* estimate of 4,434,261 individuals reached around 60 kya. However, from ∼30 kya, shortly before the LGM - the species-level trajectory diverged from the population-specific inference. Effective population size first declined gradually and then more sharply from ∼15 kya to the present. Importantly, this pattern was recovered both when singleton variants were included (**Figure 1**) and when they were excluded (**Figure S3**) from the analysis. Comparison with palaeoclimatic reconstructions from the central summer rainfall zone of Southern Africa [56] revealed that this period of *N_e_* decline coincided with repeated localised arid episodes occurring at approximately 15 – 11, 7–5 and 1.5–0.5 kya (**Figure 1**).

**Figure 1:**
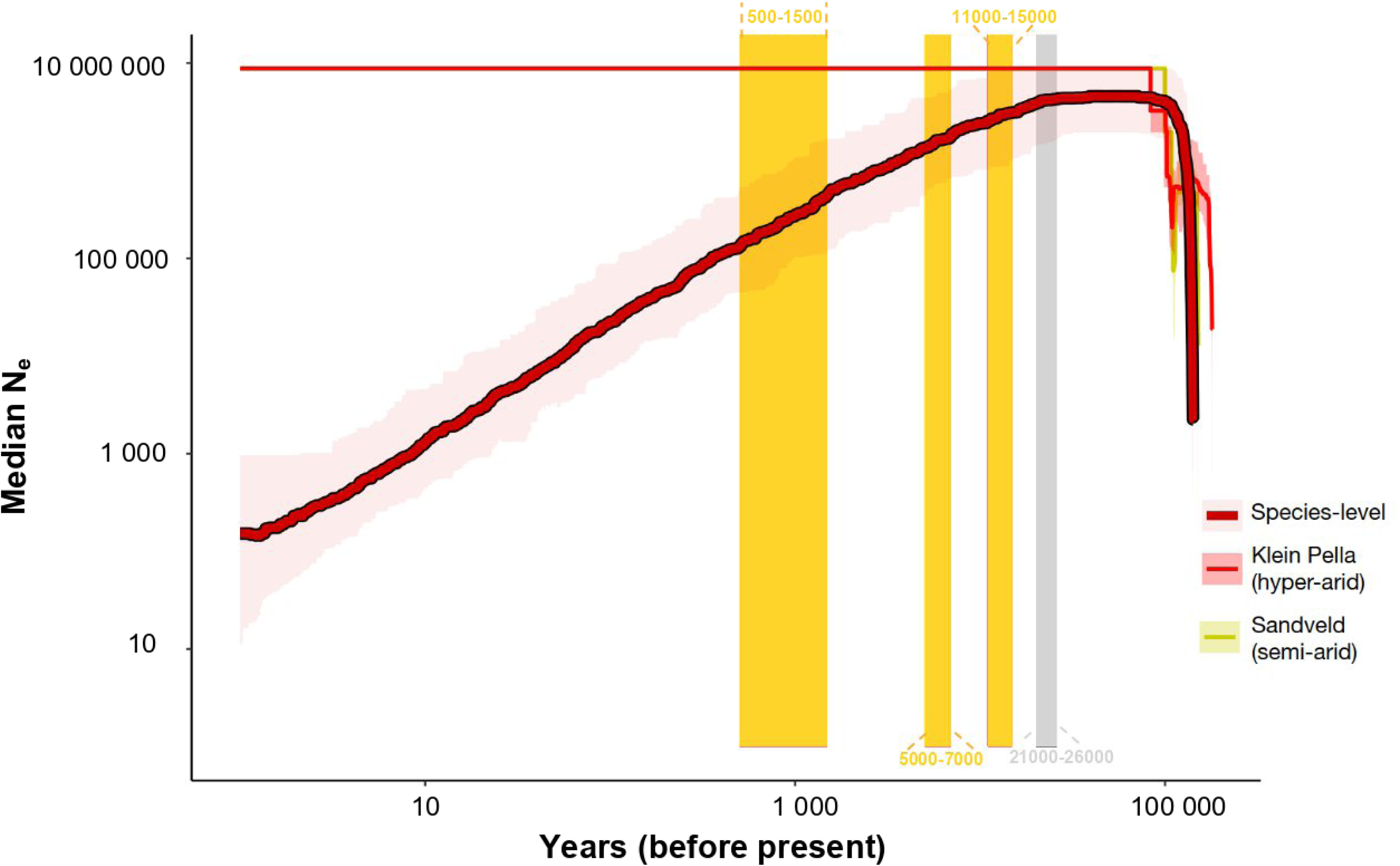
Historical changes in effective population size in *R. bechuanae*. *Stairway Plot 2* analyses (with singletons) showing historical changes in effective population size (*N_e_*) over the past 130,000 years. The thick red line indicates the estimated median *N_e_*, for the whole species (*N*=129, 146 896 SNPs), the Klein Pella locality (hyper-arid climate, *N*=14, 146 534 SNPs) is represented in thin red and the Sandveld locality (semi-arid climate, *N*=16, 145 767 SNPs) is represented in green. The lines represent the median, and the buffers represent the 95% confidence interval. The grey rectangle marks the period of the Last Glacial Maximum (LGM), while the yellow rectangles denote recent arid episodes in the Summer rainfall region of Southern Africa [63].

#### Contemporary *R. bechuanae* population structure

The analysis of contemporary population structure on all retained *R. bechuanae* individuals revealed some degree of genetic clustering despite low level of intraspecific differentiation within this species (**Figure 2**, **Tables S6-S7**). Pairwise *F_ST_* estimates among *R. bechuanae* sampling localities were relatively low, spanning from 0.010 to 0.058 (**Table S6**). Combined, the Principal Component Analysis (PCA) (**Figure S4**) and the neighbour-joining tree (**Figure 2C**) did not indicate a clear-cut separation of several sampling localities or groups of localities from each other. In contrast, the *sNMF* analysis identified three distinct genetic clusters (lowest cross-entropy value at K=3; **Figure 2**, **Table S7**). Very similar cross-entropy values for K = 3 and 4 clusters (**Figure S5**) suggest that these two inferences of population structure are almost equally likely (**Figure 2A**). At K=3, a first *R. bechuanae* genetic cluster grouped most localities from a region corresponding to the Southern Kalahari (Lake Naute, Mariental, Tswalu, Molopo, Kolomela, Sandveld), which ranged from a hyper-arid to a semi-arid climate according to the estimated Aridity Index. Another cluster was more prevalent in the Gariep and Tussen die Riviere semi-arid localities, and a third cluster was mostly found in the hyper-arid Klein Pella locality (**Figure 2B**, **Table S7**). Genetic differentiation between these groups of localities was also visible on the neighbour-joining tree (**Figure 2C**). Most individuals from localities located centrally in our sampling scheme (Benfontein, Kolomela Mine) did not have ancestry coefficients > 85 % from one specific cluster and were instead composed of mixed continuous ancestries (**Figure 2C, Table S7**), as also suggested by the PCA plot (**Figure S4**).

**Figure 2:**
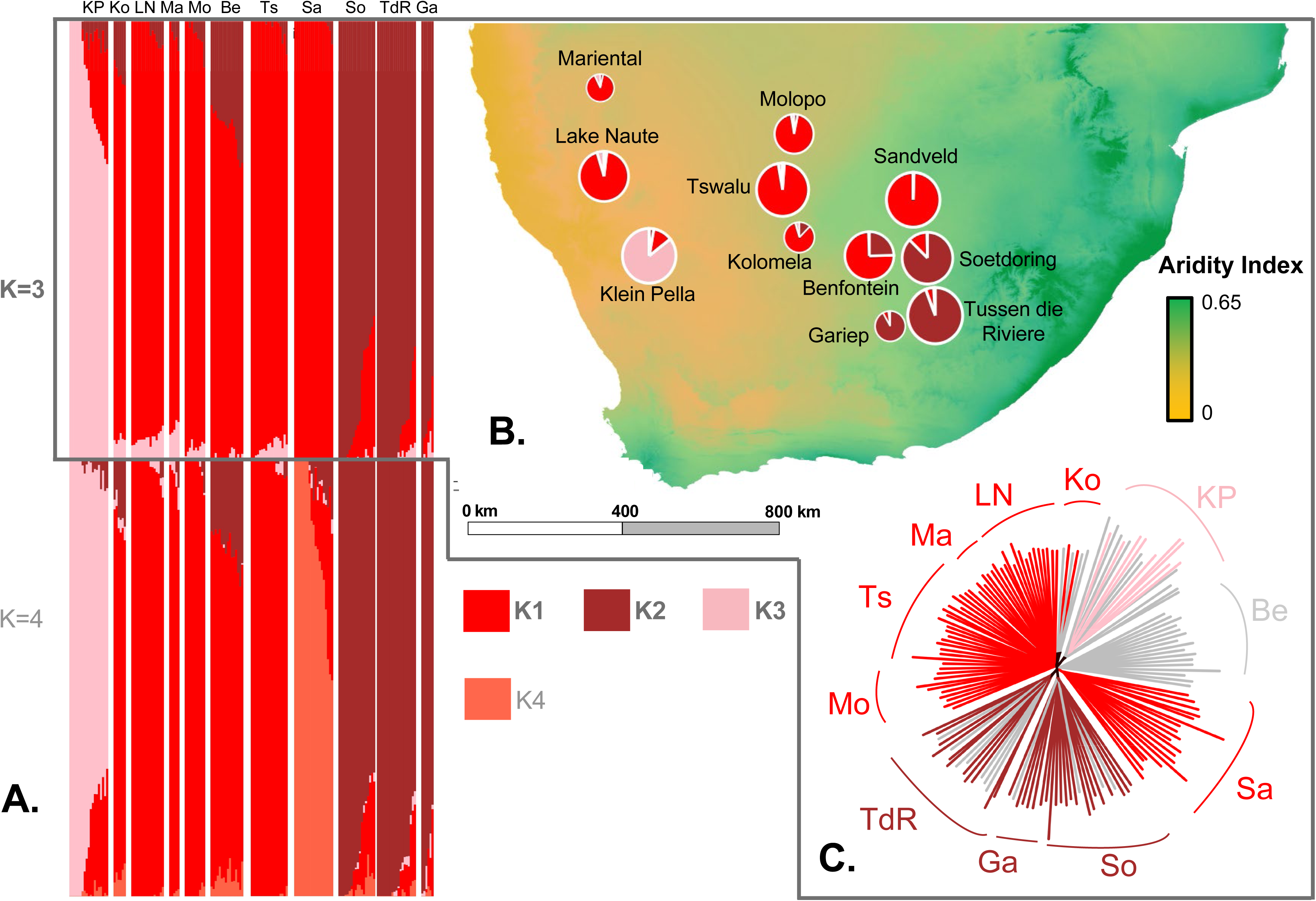
Population structure among *R. bechuanae* samples. **A**. *sNMF* results from K=3 to K=4 for *R. bechuanae* samples in each locality. Abbreviations of sampling localities are : Klein Pella (KP); Kolomela Mine (Ko); Lake Naute (LN); Mariental (Ma); Molopo (Mo); Benfontein (Be); Tswalu (Ts); Sandveld (Sa); Soetdoring (So); TdR (Tussen die Riviere); Ga (Gariep Dam). **B.** Map showing the distributions and genetic compositions of *R. bechuanae* sampling localities. Pie diagrams indicate the frequencies of genetic clusters in each locality. Base map: World Topographic Map Esri Standard, Aridity Index layer was computed from a 0.5° global grid, using data from Version 3 of the Global Aridity Index and Potential Evapotranspiration Database [35]. **C.** Neighbour-joining tree built using *ape.* Individuals with an ancestry coefficient > 0.85 for one cluster are coloured according to the colour code of the corresponding genetic cluster used in panel A. *R. bechuanae* individuals without an ancestry coefficient > 0.85 for any genetic cluster are coloured in grey. All analyses were performed using *N*=154 individuals and 44,900 SNPs.

Consistent with the gradual pattern of genetic ancestries observed across the sampled localities (**Figure 2**), simple Mantel tests using genetic distances based on *F_ST_* values among *R. bechuanae* localities (**Table S6**) showed significant positive correlations between genetic distance and geographic distance (*p* = 0.038), indicating isolation by distance, as well as between genetic distance and the environmental variable of interest, the Aridity Index (*p* = 0.037), indicating isolation by environment (**Figure 3**). Since we found a very strong correlation between geographic and environmental distances (*p* <0.001) and no significant partial Mantel tests (**Table S8**), these results suggest that patterns of isolation by distance and by environment are closely intertwined, and that genetic differentiation may be shaped by the aridity gradient, which closely mirrors the geographic distance gradient.

**Figure 3:**
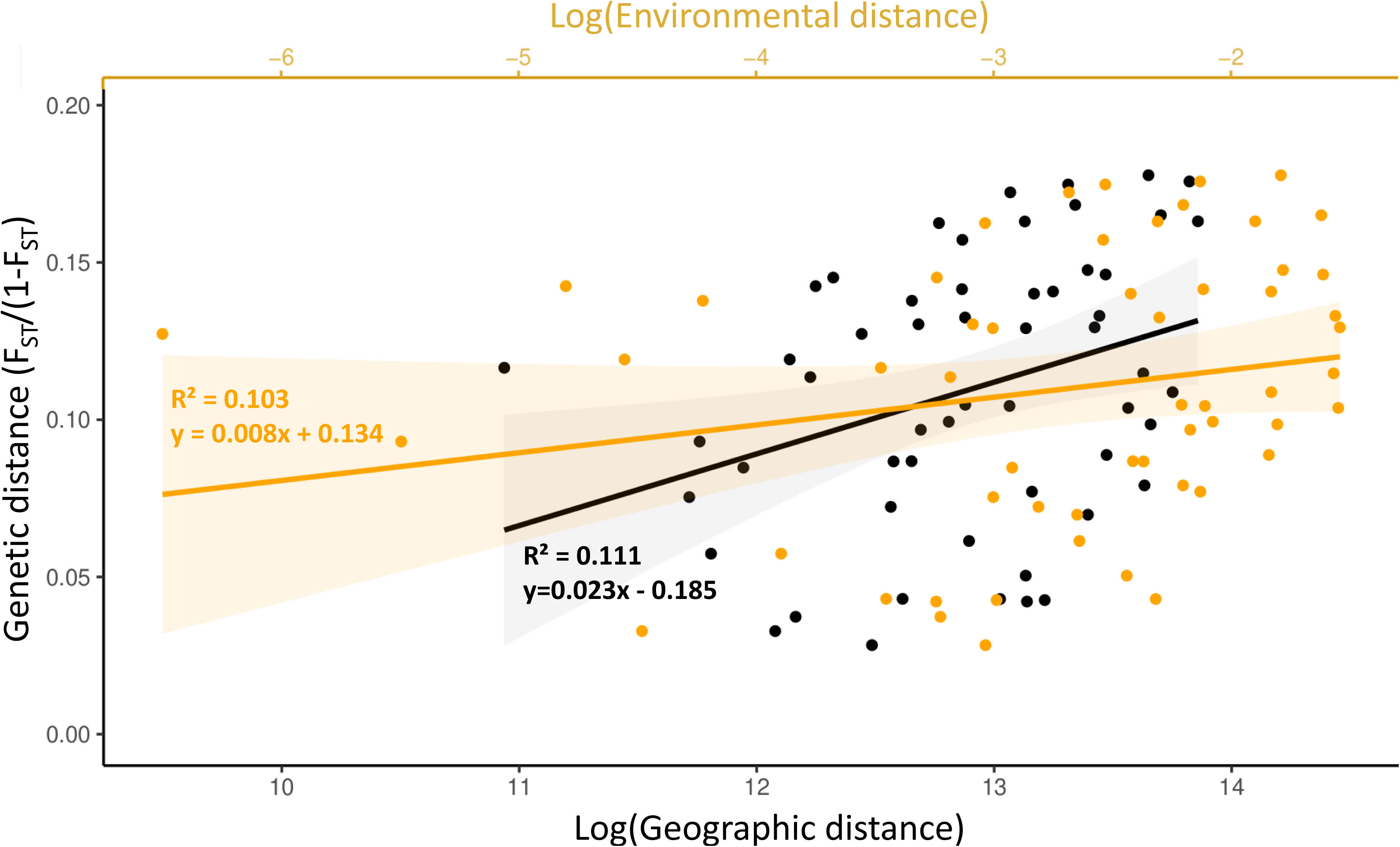
Correlations between genetic distance and geographic/Aridity Index distances. Correlation plots with slopes and correlation coefficient *R^2^* estimates between genetic distance (*F_ST_*/(1-*F_ST_*)) and log(geographic distance), and between environmental distance and geographic distance matrices (orange). Environmental distance is represented by Euclidean distance in terms of the Aridity Index.

### Genetic basis of adaptation to aridity

#### Genotype-Environment Association analysis and genomic scan for differentiation

To take advantage of the full aridity gradient across area of distribution of *R. bechuanae*, we performed the GEA analysis at the locality level using *BayPass*. Given the above correlations, we expected stringent conditions for the GEA analysis. Indeed, since *BayPass* explicitly standardises population allele frequencies for their correlation structure - here induced by the neutral population structure but also the isolation by environment signal - we expected a high rate of false negatives when testing for an association with the Aridity Index.

Overall, despite these very stringent conditions, 108 SNPs were found to be significantly associated with the Aridity Index (significance threshold of 10 dB), distributed across all genomic scaffolds (**Figure 4A**, **Table S9**). Among these, 68 were mapped within an annotated coding region of the *R. pumilio* genome, corresponding to 63 outlier ‘association’ genes in total.

**Figure 4:**
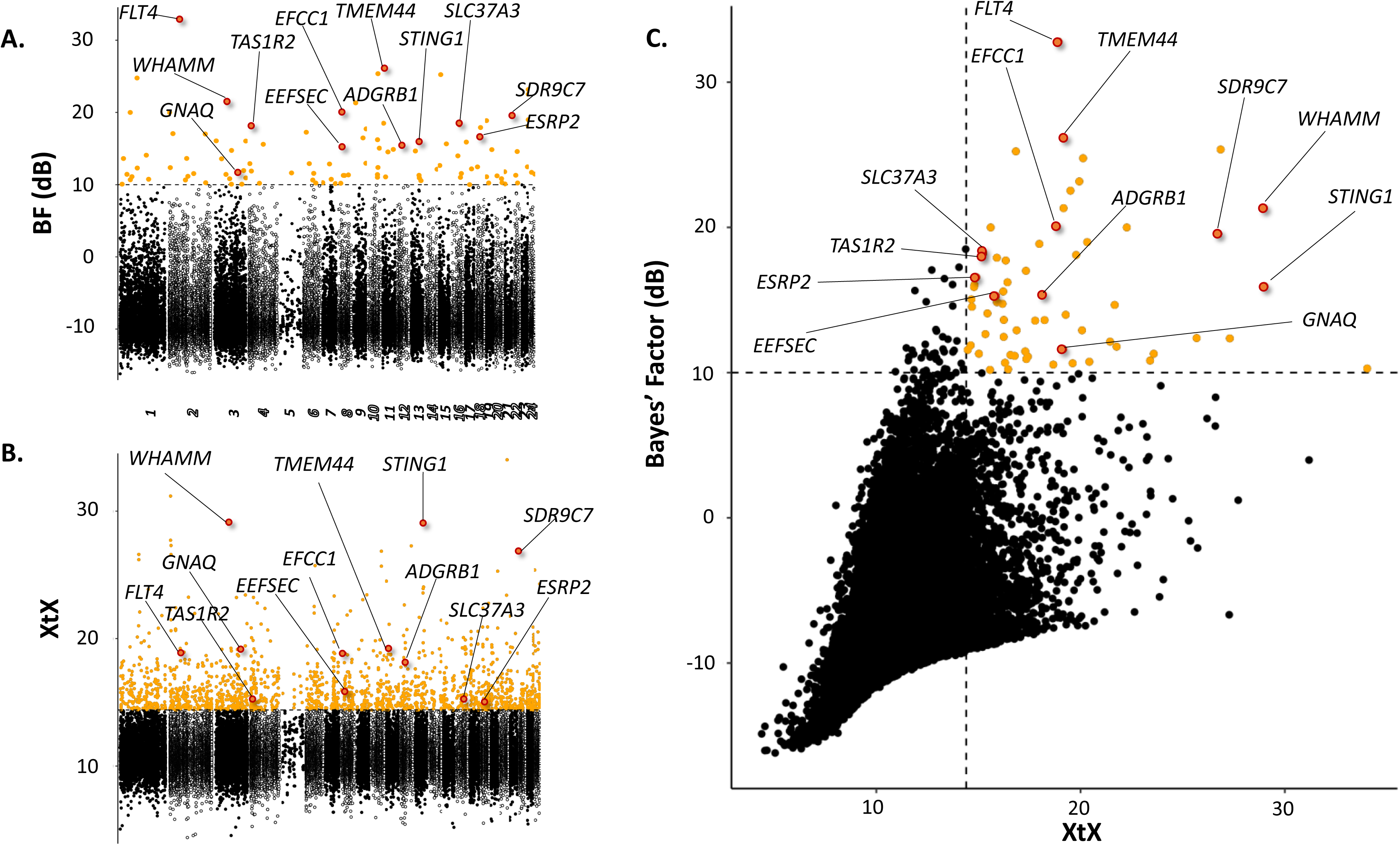
Genotype-Environment Associations and genetic differentiation in *R. bechuanae*. **A.** Manhattan plot of Bayes Factor (*BF*, in decibans) values from *BayPass* (standard covariate model), showing associations between 44,900 SNPs and the Aridity Index across 24 chromosome-level scaffolds (x-axis : genomic position). The dashed line marks the *BF*=10 (“strong” level of association) significance threshold. Outlier SNPs are in colour; others alternate black/white fill colour by scaffold). **B.** Manhattan plot of genetic differentiation (*XtX* values) from *BayPass* (core model). The dashed line indicates the 5% pseudo-observed dataset (POD) significance threshold (*XtX* = 14.4). Colour scheme as in A. **C**. Joint plot of *BF* and *XtX* values. Thresholds are shown as dashed lines; SNPs significant for both statistics are in orange. In all plots, we show the names of some genes containing significantly associated and differentiated SNPs, involved in functions relevant to the response to arid conditions (for each function, the gene containing the SNP with the highest BF was highlighted; SNPs within these genes in red). All analyses were performed using *N*=218 individuals and 44,900 SNPs.

Across the three runs of the model, which gave similar results, a total of 1,682 variants were identified as *XtX* outliers (5% *XtX* significance threshold of 14.4, see **Supplementary Material**) (**Figure 4B**). Of these, 65 SNPs displayed strong evidence of association with aridity (significance threshold of 10 dB), as well as a significant *XtX* value. Among them, 45 were mapped within an annotated coding region of the *R. pumilio* genome, corresponding to 41 outlier ‘association/differentiation’ genes.

#### Functional analyses

These 41 ‘association/differentiation’ outlier genes corresponded to twelve broad functional physiological pathways (**Table S9**), reflecting diverse metabolic functions. In **Figure 4**, we present one representative gene from each pathway, selected for showing the strongest signal of association with the Aridity Index. Among the most promising candidate genes were those involved in nervous system functions (e.g., *ADGRB1*, *WHAMM,* see **Table S9**). Notably, Gene Ontology (GO) terms related to neurotransmission showed the strongest degree of enrichment (*p_adj_*< 0.01) among genes associated with the Aridity Index, with twelve GO terms significantly enriched overall, as compared to the background genome **(Figure 5, Table S10**).

**Figure 5:**
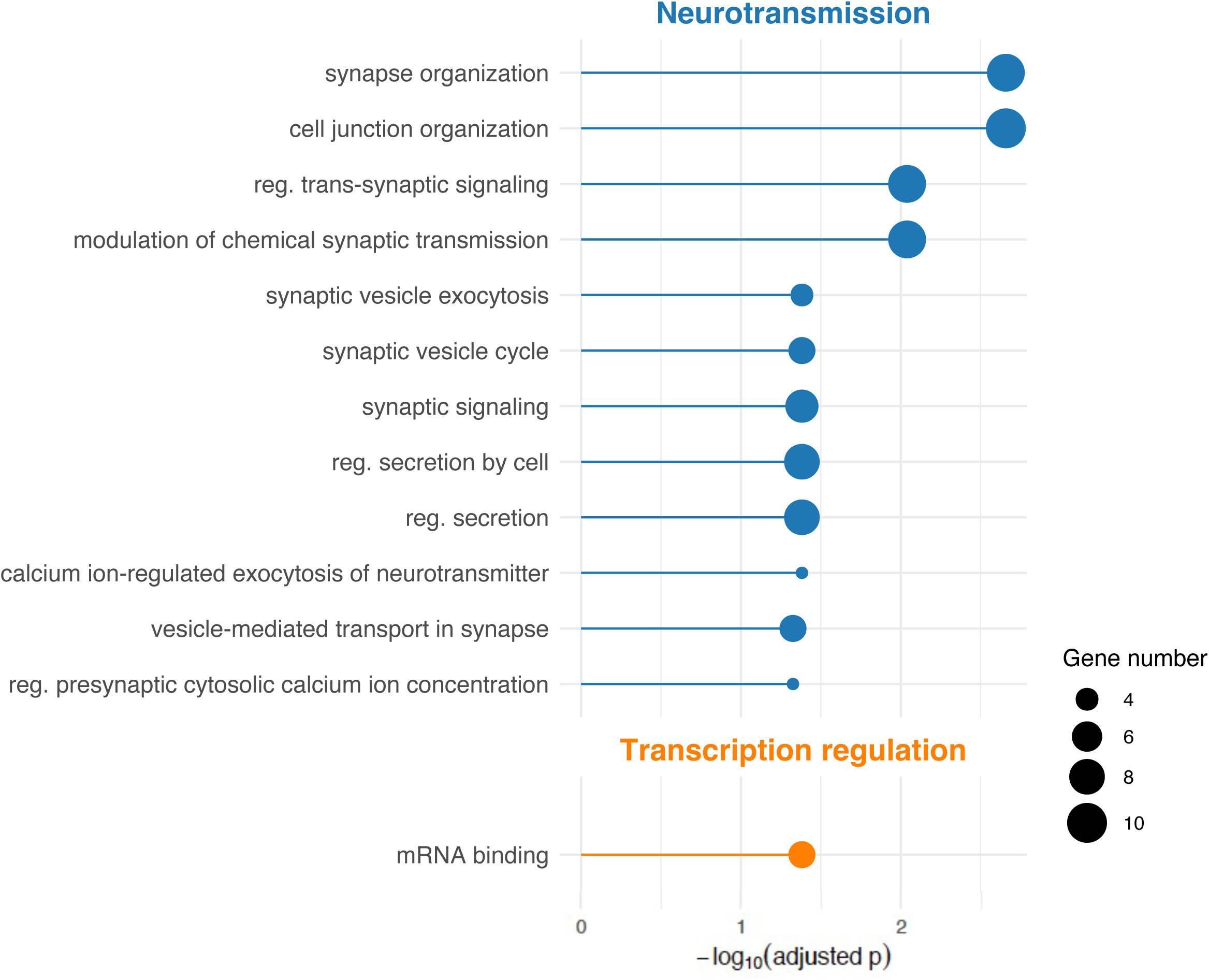
Functional enrichment among genes significantly associated with the Aridity Index. Abbreviated descriptions of significantly enriched GO terms (*p_adj_* < 0.05) are presented on the left, classified into more general physiological pathways, with level of significance after Benjamini-Hochberg correction reported on the x-axis. Circle size represents the number of outlier genes found to be associated with each GO term.

Outlier genes were also linked to maintaining osmoregulation (e.g., *EFCC1*), cellular integrity (e.g., *SLC37A3*) and stability of protein (e.g., *TMEM44*) or genetic (e.g., *ESRP2*) information, in the latter case supported by both ‘association’ and ‘association/differentiation’ outlier gene sets (**Table S9**). Relevant GO terms were significantly enriched among genes associated with aridity (**Figure 5**), and approached significance (*p_adj_* = 0.063) among associated/differentiated genes **(Table S10**). Furthermore, GO terms related to cytoskeleton regulation and GTPase activity were also near the threshold for enrichment significance in the associated/differentiated gene set (*p_adj_* = 0.063) (**Table S10**), with genes such as *WHAMM* or *EEFSEC* among top candidates (**Figure 4, Table S9**).

Finally, several genes with potential relevance for energy conservation were identified. These included genes involved in lipid metabolism (e.g., *SDR9C7*), carbohydrate metabolism (e.g., *GNAQ*), immunity (e.g., *STING1*), and sweet taste perception (*TAS1R2*, see **Table S9**).

## Discussion

In this study, we generated RAD-seq data, enabling us to investigate population-level patterns of intraspecific genetic diversity and responses to aridity in the African four-striped mouse, *Rhabdomys bechuanae*, a rodent species broadly distributed across an aridity gradient in Southern Africa. By developing an original integrative approach, combining genomic data with fine-scale contemporary and historical aridity data and leveraging a natural aridity gradient to investigate signatures of aridity-driven intraspecific divergence using GEA analyses, our study provides significant insights into population-level responses to increased aridity in a homeothermic rodent species. We implemented a strong multi-level filtering process, followed by one of the most conservative GEA methods [57]. This, along with the intertwined nature of geographic structure and environmental variation in our dataset, yielded a small, but robust set of genomic regions associated with aridity, that are further supported by meaningful functional characteristics.

Our results suggest a recent decline in effective population size at the species level, coinciding with Holocene aridification. We also show that genetic diversity within the species is shaped by variation in aridity across its distribution range. We detected allele frequency shifts associated with aridity levels, enabling identification of candidate genes and functions potentially involved in adaptation to arid environments. Although this study, sampling a portion of the genome, may miss other contributing functions, we identified candidate genomic regions associated with classical desert survival phenotypes, including DNA repair/protein modification, osmoregulation and carbohydrate/fat metabolism [10,16]. However, the strongest functional enrichment signal emerged for genes involved in neuronal transmission, bringing novel insights into the role of neurological processes in coping with osmotic stress and resource scarcity. Together, these findings support the hypothesis that genetic variation in *R. bechuanae* reflects both its demographic history, shaped by past climatic fluctuations, and the selective pressures imposed by spatial variation in aridity across its distribution range [58].

### Population structure and historical changes in effective population size in response to aridification

Although genetic differentiation amongst *R. bechuanae* localities was relatively low overall, our analyses revealed genetic clustering (three clusters) with the *sNMF* approach, along with significant correlation between genetic and geographical distances. This intraspecific structure was also supported by both PCA and neighbour-joining analyses. The observed pattern of isolation by distance provided evidence of restricted gene flow across the species’ distribution range, which aligns with the patchiness of favourable habitat available for *R. bechuanae* in the arid and semi-arid regions of Southern Africa [23,59]. Isolation by distance results from constraints on dispersal, through habitat patchiness for example, thereby promoting the population structure observed here despite the species’ relatively continuous regional distribution. Our findings support this hypothesis through the detection of significant isolation by environment associated with the Aridity Index variable, although this variable showed strong correlation with geographic distance. The population structure analysis identified genetic clusters corresponding to populations at the opposing ends of the aridity gradient (Klein Pella representing the arid extreme, and Sandveld the semi-arid extreme), both of which have relatively peripheral positions in the species’ range. This spatial arrangement suggests that the observed genetic structure could reflect adaptation to distinct environmental conditions along the aridity gradient, supporting the hypothesis that aridity is a key driver of population structure in this species.

Historical demographic events, such as range expansions, population bottlenecks or divergence due to local adaptation or neutral processes, may have contributed to the contemporary patterns of genetic diversity and population structure in *R. bechuanae.* To investigate whether such changes were associated with past climatic events, we inferred historical changes in effective population size (*N_e_*) and compared them with available palaeoclimatic reconstructions.

SFS-based demographic inference from reduced-representation datasets is sensitive to data quality and to the representation of rare variants. Although our filtering strategy aimed to maximise data quality while retaining rare alleles, the species-level SFS showed a deficit of singleton variants relative to neutral expectations. This pattern most likely reflects the loss of true singletons during stringent filtering to reduce genotyping errors and missing data. Similar singleton depletion has been reported in RADseq and low-coverage datasets, where filtering disproportionately removes ultra-rare alleles ([60–61]. Because rare alleles disproportionately inform recent coalescent events, demographic inference near the present is particularly sensitive to singleton accuracy and sequencing artefacts (e.g, [62]). The most recent portion of inferred trajectories should therefore be interpreted with caution. Importantly, our results are robust to these limitations. Removing singleton variants did not alter the inferred demographic trajectories, indicating that the main signal is supported by multiple allele-frequency classes.

At the locality level, a reduction in *N_e_* was detected in the hyper-arid Klein Pella population around the Last Glacial Maximum (LGM), consistent with increased aridity and climatic instability in southern Africa during this period [56,63]. In contrast, both localities showed stable *N_e_* from the LGM to the present, with no evidence of strong recent decline. At the species level, a decline in *N_e_* beginning ∼15 kya was inferred. This signal was robust to the exclusion of singletons, suggesting that it is not driven by rare-allele artefacts. However, because SFS-based inference assumes panmixia and this pattern was not recovered in locality-level analyses, the inferred species-level decline may primarily reflect population structure and changes in connectivity within *R. bechuanae* rather than true changes in census size. Indeed, simulations have shown that apparent signals of past changes in population size can often arise from stable population structure or changes in population connectivity [64–66]. The observed pattern may therefore reflect a complex combination of intra- and inter-deme processes that are difficult to disentangle using SFS-based demographic approaches [64]. Nevertheless, given the coincidence between the inferred decline in *Ne* and repeated localised arid episodes [56], changes in population structure and connectivity may themselves have been driven by fluctuations in aridity. Recurrent arid phases during the Late Pleistocene and Holocene likely fragmented habitat, by altering vegetation and nesting site availability, thereby reducing gene flow among populations [56,67,68], and potentially producing the observed pattern.

Overall, our results support the hypothesis that past climatic fluctuations in the Southern Kalahari and central South Africa may have been important factors shaping effective population size and population structure of *R. bechuanae,* although other neutral and adaptive factors could also be involved. Historical periods of increased aridity could not only have reduced population connectivity but may have also created opportunities for local adaptation if geographically separated populations experienced different environmental pressures along the aridity gradient. This dual effect of aridity, both fragmenting populations and creating divergent selective environments, provides a comprehensive explanation for the observed genetic structure in *R. bechuanae*.

### Adaptive genomic variation along an aridity gradient

This study enabled the identification of several SNPs associated with the Aridity Index variable, despite the very stringent conditions of the *BayPass* analysis due to significant correlations between genetic, geographic and environmental (Aridity Index) distances. While these conditions reduced the overall power of the analysis, they also strengthened the reliability of the identified SNPs. Additional approaches, such as genome-wide association studies based on well characterised physiological phenotypes [28], could complement our findings by revealing other genomic regions underlying key traits for adaptation to aridity. It is unlikely our inferences were biased by small sample sizes in some localities, as GEA studies using Bayes’ factor are not particularly sensitive to sample sizes [69]. We only focused on SNPs mapping inside the coding region of genes, hence proposing thus providing a very stringent view of the genes associated with the Aridity Index; more genome regions, in linkage disequilibrium with the regions we identified, might also contribute to the response to aridity, but we decided to focus on the most robust genomic regions to interpret our results functionally.

Our hypothesis of differential selective pressures acting on *R. bechuanae* populations along the aridity gradient was supported by the identification of outlier SNPs located within coding regions of genes associated with important biological functions for adaptation to an arid or desert lifestyle. We identified genes involved in energetic balance (e.g., lipid metabolism, immunity) and osmotic balance (e.g., osmoregulation, neuronal processes, body fluids regulation), as well as cellular responses to dehydration (e.g., protein metabolism, regulation of transcription and cytoskeleton). *Rhabdomys bechuanae* faces environmental constraints, including high temperatures and resource scarcity. Even in semi-arid conditions, these challenges cause a strain on homeostasis, as revealed in *R. bechuanae* populations from semi-arid locations by blood markers of osmoregulation, immunity and energy levels [28]. More arid conditions could cause cellular stress, and exacerbate the effects of dehydration due to lack of water and rapid water loss (e.g., respiratory water loss, evaporative cooling). Overall, consistent with our predictions, our results suggest that various physiological mechanisms may enable *R. bechuanae* to cope with harsh conditions and inhabit drier environments. We synthesised previous findings from other species aligning with our results in **Table S9**, and discuss in more detail the functions of interest for adaptation to arid conditions in light of this literature review.

In numerous studies carried out on mammalian species, the same genes, or those involved in similar functions as reported in our study, are highlighted as containing SNPs varying with an aridity climatic index (e.g., Moisture index [70]), differentially expressed in arid conditions (e.g., water restriction [19]), or showing signs of selection in specialist organisms (e.g. [15]). This intersection suggests a genetic basis for adaptation to aridity partly shared across various mammals. Studies on the basis of adaptation to xeric environments show convergent changes in tissue composition, gene expression and coding sequences, with significant overlap mostly among related species [10,71].

Our findings align with key shared mechanisms of adaptation to low resource availability observed across mammals, especially those involved in maintaining energetic balance, such as lipid metabolism [10,13], and immune response [72]. Interestingly, acquiring energy in a food-scarce environment may also involve diet shifts in response to arid food sources, such as desert plants and insects. These shifts can be facilitated by mutations in taste receptor genes, such as the sweet taste receptor identified in our study or in the numbat *Myrmecobius fasciatus* [73], and bitter taste receptors in the *Peromyscus eremicus* cactus mouse [14].

At the cellular level, dehydration-induced changes, such as cytoskeletal rearrangements [74], may explain the observed association between aridity and genes regulating the cytoskeleton (e.g., GO terms: filopodium assembly), suggesting possible adaptations to hyperosmotic stress in the striped mouse. However, dehydration also affects cells through protein denaturation, transcription/translation suppression, altered adhesion, increased DNA breaks and protein oxidation, cell cycle arrest, and apoptosis [74–76]. Consistent with these responses, SNPs associated with aridity were found in genes involved in protein synthesis and modification, DNA repair, and cellular cycle regulation. Genes containing outlier SNPs were also enriched for GO terms related to transcription regulation. These findings align with previous results from selective sweeps studies in arid-adapted rodents (e.g., *Peromyscus eremicus* [14]), and meta-analyses of metazoan genomic and transcriptomic data showing selection on translation-related genes in harsh environments [77]. Indeed, cellular responses to hyperosmotic stress commonly involve regulating protein translation and degradation [76]. Overall, our results suggest strong selection on genes mitigating thermal or hyperosmotic stress and restoring cellular function after acute stress [75].

Arid environments impose strong selective pressures on osmoregulation mechanisms. In *R. bechuanae*, we identified genes containing significantly associated outlier SNPs involved in osmoregulation, many of which were also reported in other desert rodents (e.g., *Allactaga sibirica*, *Dipus sagitta*, *Meriones meridianus* and *Phodopus roborovskii* [15], **Table S9**). Water balance in mammals is regulated by brain-body communication mediated by central nervous system structures that exhibit marked plasticity in response to environmental cues in arid-adapted species [78]. This may help explain the strong enrichment of GO terms related to nervous system functions, such as nervous system development and neurotransmitter regulation, which emerged as the most significant signal in our study. Although neurological functions are occasionally noted in aridity adaptation studies (e.g., [13,70,79]), they are rarely emphasised as key survival factors under arid conditions. When mentioned, they are typically discussed in the context of selective brain cooling in large, heat-adapted mammals [80]. Neurotransmitters, essential for maintaining homeostasis [81], regulate blood flow, a critical function in arid environments where dehydration can restrict organ perfusion [82]. Our identification of outlier genes involved in heart rate regulation and vasculature development - also differentially expressed in other desert-adapted rodents such as *Abrothrix olivacea* [83], supports this. Together, these results support the involvement of a previously underappreciated functional category – neurological pathways - in the genomic response to aridity, offering new directions for research in the evolutionary physiology of aridity adaptation.

## Conclusion

This study sheds new light on how historical climatic fluctuations and environmental gradients shaped population structure and genetically based responses to aridity in *Rhabdomys bechuanae*, a rodent species inhabiting some of the most arid regions of Southern Africa. By integrating genomic data with fine-scale environmental information, we uncovered signatures of population differentiation and polygenic adaptation along an aridity gradient, involving functions linked to osmoregulation, energetic balance, and cellular stress responses. The list of candidate genes and biological functions provides a valuable foundation for further research on the molecular mechanisms of adaptation to arid environments. While we focused on coding regions to identify genes and functions associated with the Aridity Index, future work incorporating transcriptomic and non-coding genomic data will be essential to assess the role of gene expression regulation in this species’ response to increasing aridity and to disentangle the relative contributions of coding versus regulatory variation. Moreover, although this study revealed genetically based signatures of adaptation, avoidance behaviours and plastic physiological responses likely play a critical role in coping with environmental stress imposed by seasonal variation in aridity [79]. Future studies manipulating water and food availability, will be key to elucidating the interplay between phenotypic plasticity and local adaptation in shaping species’ responses to arid conditions.

