## Supplementary Material for "Role of aridity in shaping adaptive genomic divergence and population connectivity in a Southern African rodent"

**Supplementary Methods**

RAD-seq protocol

DNA extractions were performed from tail tissues preserved in 98% ethanol, using a 96-Well Plate Animal Genomic DNA Miniprep Kit (BioBasic Inc.). Since samples from other *Rhabdomys* species and subspecies had also been collected for a separate study, we proceeded with RAD-seq library preparation and sequencing for *R. bechuanae* samples analysed in this study concurrently with the other *Rhabdomys* samples (*N* = 411 in total). Library preparation followed the protocol of Etter et al. [1], with modifications as described in [2]. DNAs were digested with the 8-cutter restriction enzyme *SbfI* and ligated to a modified Illumina P1 adapter containing individual multiplex identifiers (MIDs) 5-6 bp long. Then, 20 pools of libraries were prepared based on equimolar mixes of 22 to 29 digested and tagged DNAs per pool (global *Rhabdomys* sample set including 230 *R. bechuanae* individuals). Each pool was sheared by sonication using an S220 ultra-sonicator (Covaris, Inc.) and size-selected for 300–600 bp by agarose gel excision. Different P2 adapters including pool multiplex identifiers (MIDs) were then ligated to different pools intended to be sequenced on the same sequencing lane. The DNA fragments containing both adapters (P1 and P2) were PCR enriched during 18 cycles. These 20 pools of RAD libraries were sequenced on a total of four S1 lanes and two SP lanes of an Illumina NovaSeq 6000, using a 2x150 bp paired-end protocol, at MGX-Montpellier GenomiX Facility (Montpellier, France). The reference-based *Gstacks* pipeline was applied to cluster reads from all available samples (*N* = 411) into RAD loci before using the *populations* program on *R. bechuanae* samples only to proceed to SNP and sample filtering steps.

Aridity Index

We computed the Aridity Index (AI) to estimate aridity levels of each *R.bechuanae* sampling locality. AI is typically defined as the ratio between precipitation (*P*) and potential evapotranspiration (*PET*) in a given environment [3]. For each *R. bechuanae* locality, the Aridity Index (AI) was retrieved based on data from Version 3 of the Global Aridity Index and Potential Evapotranspiration Database [4], computed on a 0.5° global grid using the 30-year average of *P* / *PET* (1981-2010)

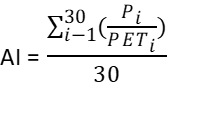

where *i* denotes the *i*^th^ year.

SNP filtering for each dataset

Dataset (A) was for population structure analyses/genetic differentiation estimates, dataset (B) for inferences of historical changes in effective population size and dataset (C) for GEA analyses. Some SNP and sample filtering steps were applied similarly to all three datasets: a minimum Phred genotype quality score of 40, a minimum read depth per genotype of three, a minimum 10X and maximum 30X coverage (based on empirical distributions), and a minimum percentage of the samples genotyped of 80 %. We also retained in each dataset only the first SNP of each locus to minimise linkage disequilibrium bias in downstream analyses, and only individuals with less than 50% of missing RAD loci. The minor variant count filter (--mac) was only applied (mac=3) to datasets A and C to minimise sequencing errors. This filter was not applied to dataset B to avoid removing large numbers of rare alleles, particularly singletons, informative for demographic inferences. Finally, a downsampling procedure was applied to datasets A and B. Unbalanced sample sizes can induce some bias in the inference of population structure [5] and demographic history [6], notable through overestimating admixture with the *sNMF* method [7]. We downsampled each locality up to a maximum of 19 samples, with only the samples with the least missing data being included for each locality, as missing data can lead to biases in population genetic inferences [8,9]. Several different parameter sets were tested for each filter parameter, partly based on empirical distributions, but intermediate stringency was ultimately chosen as a compromise between SNP numbers *vs*. coverage. The sequence order of the different SNP and sample filtering steps, their application to each analytical dataset and their effect on the number of final samples and SNP retained are provided in **Table S2**. Because site frequency spectrum (SFS)-based demographic inference is particulartly sensitive to rare alleles, especially singletons and doubletons, and missing data. we used the SNP dataset obtained without applying a minor allele count (-mac) filter and we minimised the effects of missing data while retaining a sufficient number of SNPs for inferences of historical effective population size by implementing two complementary filtering steps specific to the *Stairway Plot 2* analyses. These steps are described in the main text and data quality metrics for the final datasets used for *Stairway Plot2* analyses reported in Table **S3**.

Stairway plot inference

Ultimately, 67 % of sites were used to create an SFS training set and train the model, with the remaining sites used to create an SFS testing set for the trained model. Goodness-of-fit of the trained models was tested using ¼, ½, ¾ or n-2 “breakpoints” (defining the boundaries of each epoch), and the best-fit model was used to produce the final inference. Two hundred simulations were carried out for each estimation.

XtX outlier detection

*P*-values associated with the *XtX* calibrated estimator were not distributed as expected under the null hypothesis (**Figure S1**). Therefore, *p*-value thresholds were not used to identify outliers. Instead, *XtX* statistics were calibrated through the simulation and analysis of a pseudo-observed dataset as described in [10], simulated with neutral 44,900 SNPs, which inferred a 5% *XtX* significance threshold of 14.4.

**List of supplementary tables and figures :**

**Table S1**: Sample list and selection for downstream analyses

**Table S2** : SNP data filtering information for each type of downstream analyses

**Table S3** : Data filtering steps and quality metrics specific to SFS-based Stairway Plot2 analyses

**Table S4**: RADseq read and individual filtering results

**Table S5**: Estimates of the Aridity Index and genetic diversity for R.bechuanae sampled localities

**Table S6** : Genetic differenciation estimates (*F_ST_*) among R. bechuanae sampled localities

**Table S7** : Ancestry coefficients for each sampled individual obtained from LEA analysis (best run, K=3) across normalised *Rhabdomys bechuanae* samples

**Table S8**: Simple and partial Mantel test results

**Table S9**: List of outlier SNPs significantly associated with the Aridity Index and related information

**Table S10**: Functional enrichment results after *BayPass* Genotype-Environment Association analysis

**Supplementary tables are provided as separate Excel files or Word files.**

**Figure S1:** *XtX* *p*-value diagnostic plots

**Figure S2:** Site-frequency spectra obtained with easySFS**.**

**Figure S3:** *Stairway Plot 2* estimation of changes in effective population size (*N_e_*) through relative time in the analysis performed without singletons

**Figure S4:** Principal Component Analysis

**Figure S5:** Plot of mean cross-entropy estimates obtained with the *sNMF* analysis

**Supplementary figures are provided in this document.**

**Supplementary Figures**

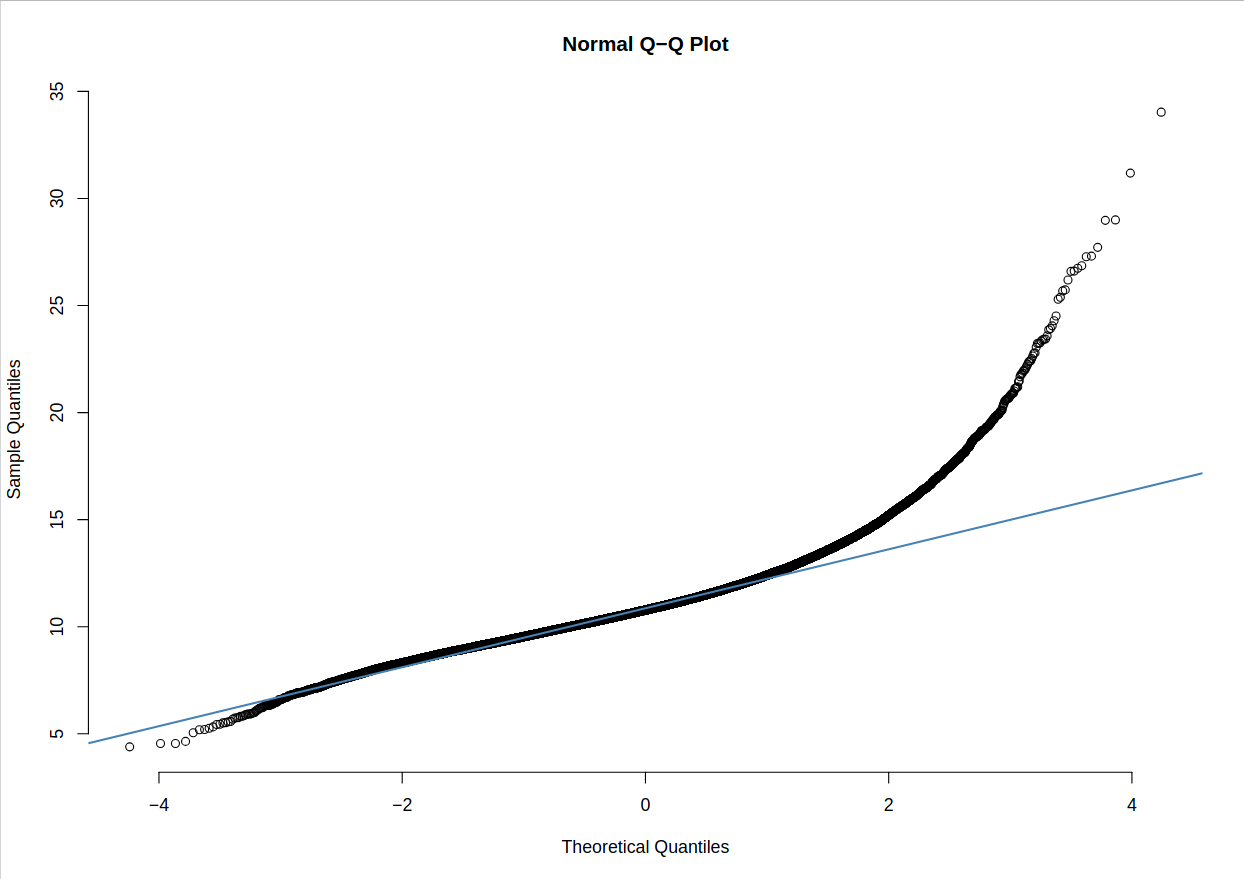

**Figure S1: *XtX* *p*-value diagnostic plots.** The residuals of the *XtX* p-values outputted by *BayPass* did not follow a normal distribution.

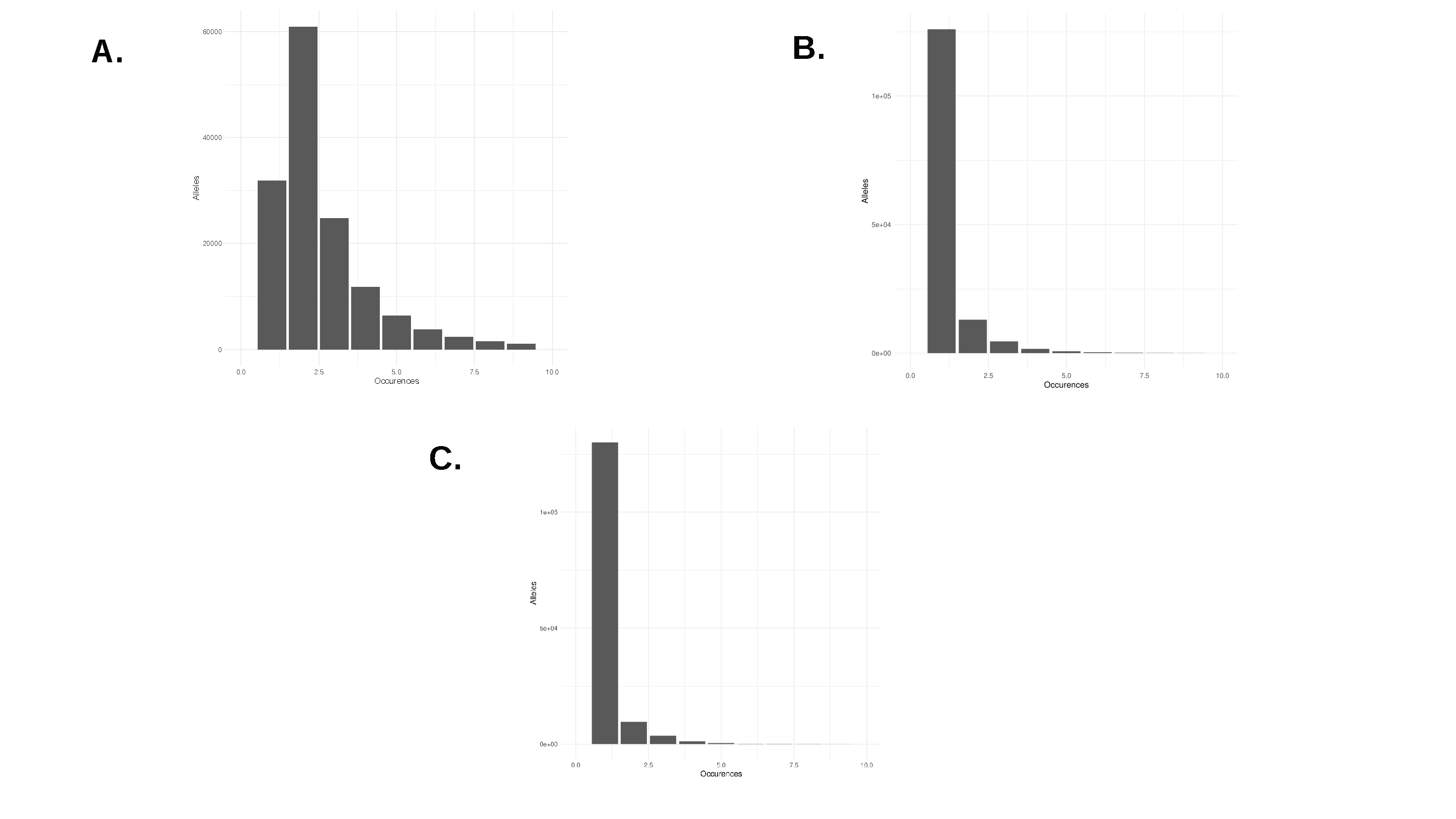

**Figure S2: Site-frequency spectra (SFS) obtained using *easySFS*.** A: species-level dataset; B: Klein Pella dataset; C: Sandveld dataset.

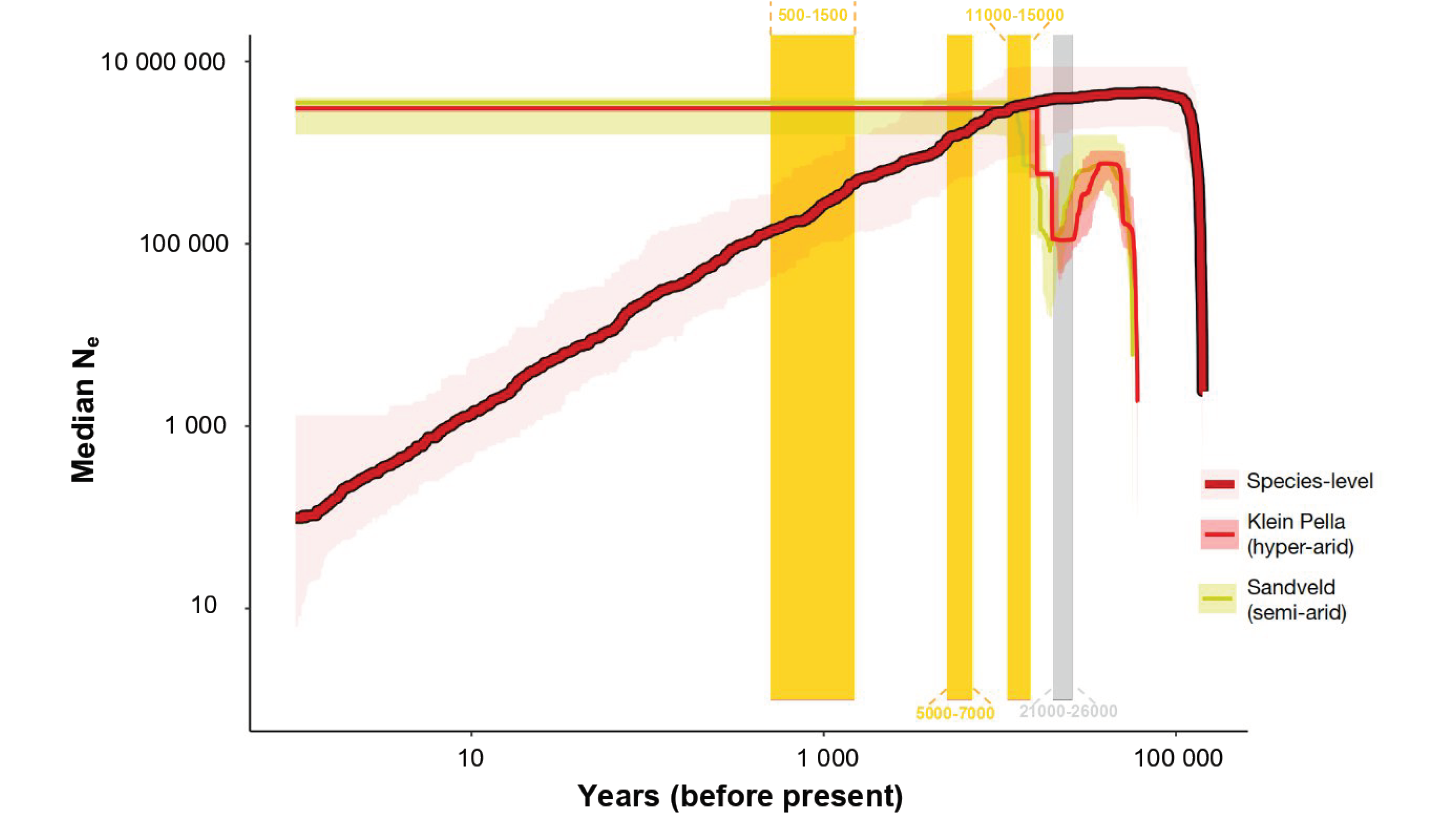

**Figure S3: *Stairway Plot 2* estimation of changes in effective population size (*N_e_*) through relative time in the analysis performed without singletons** – The lines represent the medians, and the buffers represent the 2.5 and 97.5 percentiles.

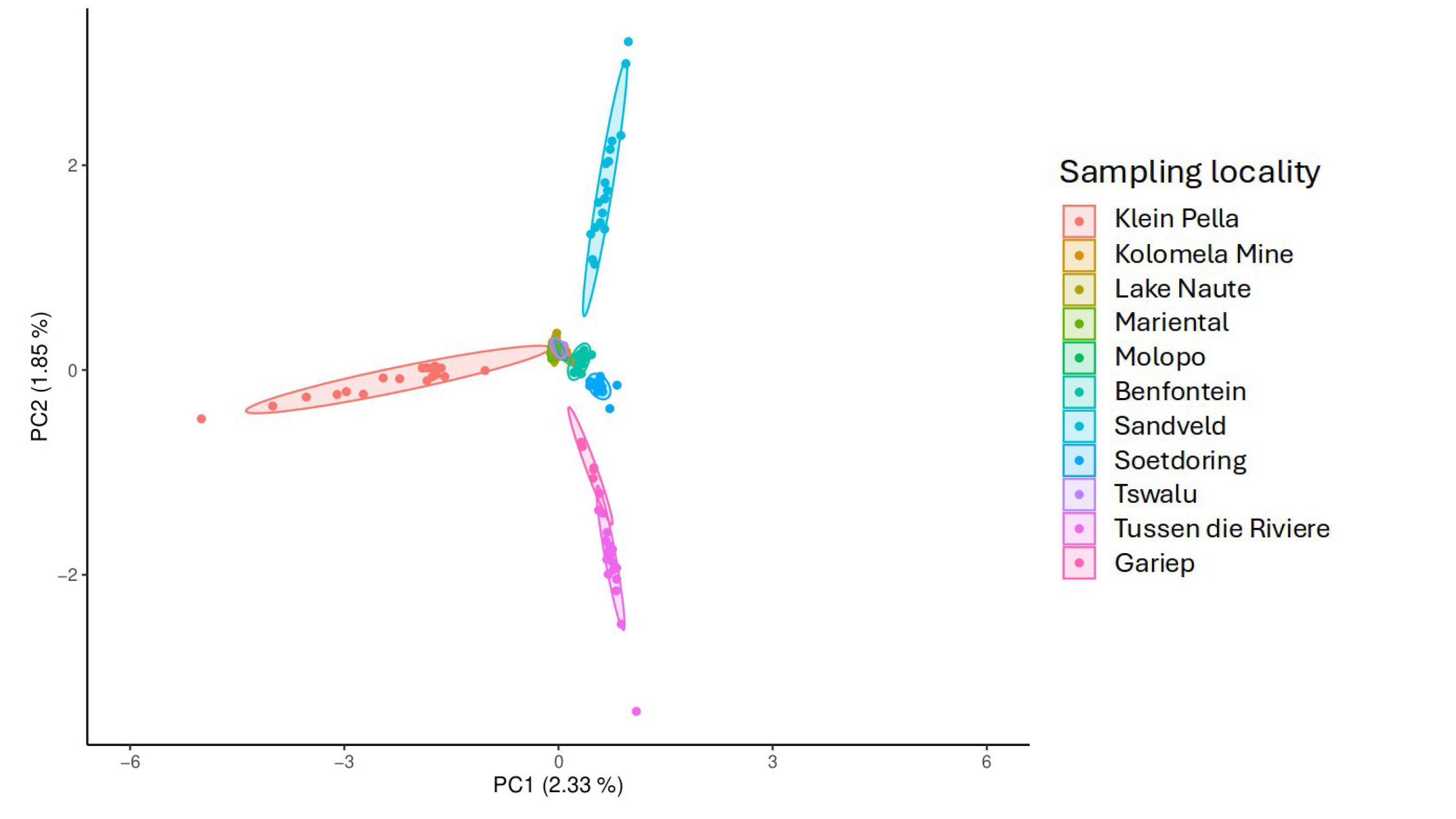

**Figure S4: Principal Component Analysis.** This plot shows the first two Principal Components of the genetic relationship matrix for the eleven sampled localities**.**

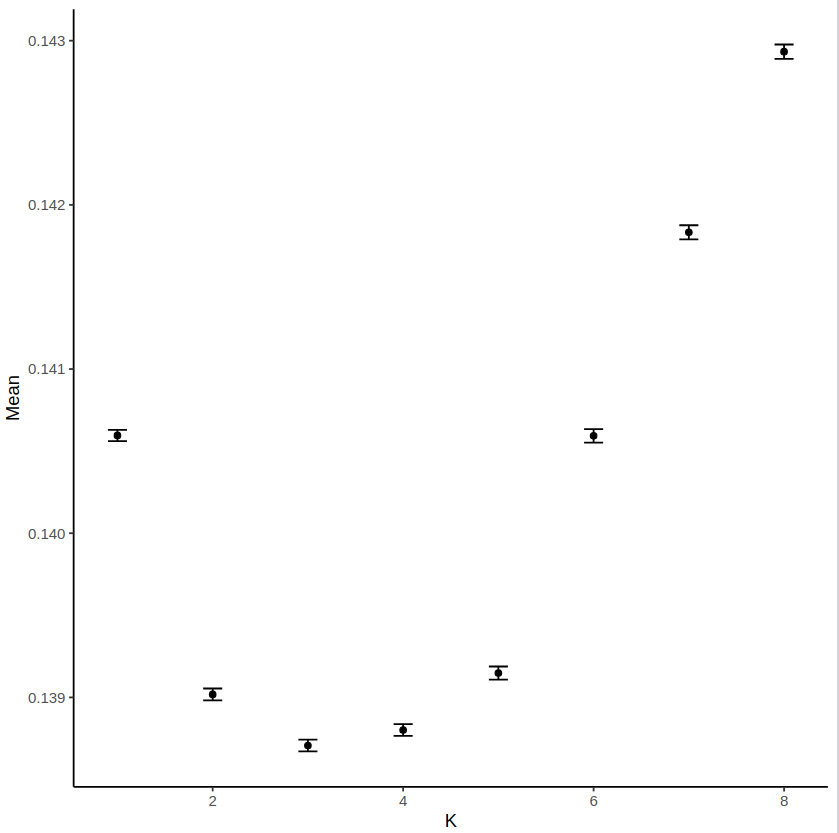

**Figure S5: Plot of mean cross-entropy estimates obtained with the *sNMF* analysis.** Estimates were obtained for varying number (1 to 8) of ancestral populations (K) in 1500 runs of sparse non-negative matrix factorisation**.**

*References*

1. Etter PD, Bassham S, Hohenlohe PA, Johnson EA, Cresko WA. SNP discovery and genotyping for evolutionary genetics using RAD sequencing. In: Methods in Molecular Biology. Totowa, NJ: Humana Press; 2012. p. 157–78. (Methods in molecular biology (Clifton, N.J.)). (https://doi.org/10.1007/978-1-61779-228-1_9)

2. Cruaud A, Gautier M, Galan M, Foucaud J, Sauné L, Genson G, et al. Empirical assessment of RAD sequencing for interspecific phylogeny. Mol Biol Evol. 2014 May;31(5):1272–4. (https://doi.org/10.1093/molbev/msu063)

3. Middleton NJ, Thomas DSG. World Atlas of Desertification. Edward Arnold. 1992.
